# CRISPRAnalyzeR: Interactive analysis, annotation and documentation of pooled CRISPR screens

**DOI:** 10.1101/109967

**Authors:** Jan Winter, Marc Schwering, Oliver Pelz, Benedikt Rauscher, Tianzuo Zhan, Florian Heigwer, Michael Boutros

## Abstract

Pooled CRISPR/Cas9 screens are a powerful and versatile tool for the systematic investigation of cellular processes in a variety of organisms. Such screens generate large amounts of data that present a new challenge to analyze and interpret. Here, we developed a web application to analyze, document and explore pooled CRISR/Cas9 screens using a unified single workflow. The end-to-end analysis pipeline features eight different hit calling strategies based on state-of-the-art methods, including DESeq2, MAGeCK, edgeR, sgRSEA, Z-Ratio, Mann-Whitney test, ScreenBEAM and BAGEL. Results can be compared with interactive visualizations and data tables. CRISPRAnalyzeR integrates meta-information from 26 external data resources, providing a wide array of options for the annotation and documentation of screens. The application was developed with user experience in mind, requiring no previous knowledge in bioinformatics. All modern operating systems are supported.

**Availability and online documentation:** The source code, a pre-configured docker application, sample data and a documentation can be found on our GitHub page (http://www.github.com/boutroslab/CRISPRAnalyzeR). A tutorial video can be found at http://www.crispr-analyzer.org.

## Introduction

In vertebrate cells, genome editing mediated by CRISPR/Cas9 can be leveraged for high-throughput screens in a pooled format (Zhou et al., 2014; Steinhart *et al.*, 2016; Wang *et al.*, 2015; Hart *et al.*, 2015; Shi *et al.*, 2015). In such screens, a lentivirus-based short guide RNA (sgRNA) library is used to infect Cas9-expressing cells, followed by a positive selection for transduced cells. Next generation sequencing (NGS) is employed to identify the enrichment or depletion of mutant cell clones based on the representation of sgRNA barcodes in the cell populations (Shalem *et al.*, 2014).

End-to-end analysis of a pooled CRISPR/Cas9 screen consists of multiple steps: First, sequencing quality and screen quality have to be evaluated, requiring handling of large amounts of NGS data. After data normalization, the optimal strategy to identify hits has to be chosen. Visualizing and annotating the results with meta-information such as Gene Ontology terms or other resources are key steps to gain new biological insights. Finally, documentation of both the analysis procedure and the screen for long term archiving ensure reproducibility and good scientific practice. Several analysis algorithms and workflows have been developed to identify interesting candidate genes in a screen.

Often, these were designed with specific selection criteria in mind, such as gene essentiality (Hart and Moffat, 2016; Yu *et al.*, 2016), drug resistance, or negative viability screens (Li *et al.*, 2014; Dai *et al.*, 2014; Diaz *et al.*, 2014; Yu *et al.*, 2016; Noh and Beibei, 2015). Comparing the results of multiple candidate calling strategies in the context of the analysis of a single screen helps to choose the most robust candidates. Currently available end-to-end pipelines only cover subsets of these steps (Winter *et al.*, 2015; Diaz *et al.*, 2014; Li *et al.*, 2015) and require additional tools to complete the screen analysis.

To fill this gap, we developed CRISPRAnalyzeR, which is the first end-to-end analysis platform for pooled CRISPR/Cas9 screens. Its web-based interface enables users to perform exploratory data analysis, annotation and documentation of a screen in a single, comprehensive workflow. Tools for the assessment of sequencing data quality and the evaluation of the screen performance help users judge the quality of their experiment. CRISPRAnalyzeR puts eight different hit-calling strategies based on state-of-the-art algorithms at the user’s fingertips, enabling them to identify robust candidates for follow up experiments. Results can be annotated with information from 26 external data resources. CRISPRAnalyzeR automatically documents all analysis steps. An interactive report can be generated, ensuring reproducibility and good scientific practice. It offers quality assessment of sequencing data, NGS data handling, the evaluation of the screen performance, parallel data analysis using eight different state-of-the-art hit calling algorithms, as well as extensive annotation using 26 external data resources and comprehensive documentation.

CRISPRAnalyzeR is an open-source tool based on R and can be used online or installed locally in a few minutes on many operating systems using a pre-configured application.

## CRISPRAnalyzeR offers comprehensive web-based analysis workflow

CRISPRAnalyzeR is designed with user experience in mind, incorporating all steps required for the comprehensive analysis and documentation of pooled CRISPR/Cas9 screens into a single application. The analysis workflow (Figure 1A) is consists of six different steps: 1) uploading the data and setup of the screening parameters (Setup Screen), 2) assessment of both the sequencing quality and the screen performance (Screen Quality), 3) identification of candidate genes using eight different algorithms simultaneously (Hit Calling), 4) annotation of candidate genes (Hit Confirmation) and 5) offline screening and analysis documentation in an interactive HTML file (Documentation).

All steps are performed via a web-based user interface as illustrated in Figure 1B. Many interactive plots and tables guide the analysis. They can be directly saved in various formats such as JPG, PDF, XLSX or CSV.

**Figure 1:**
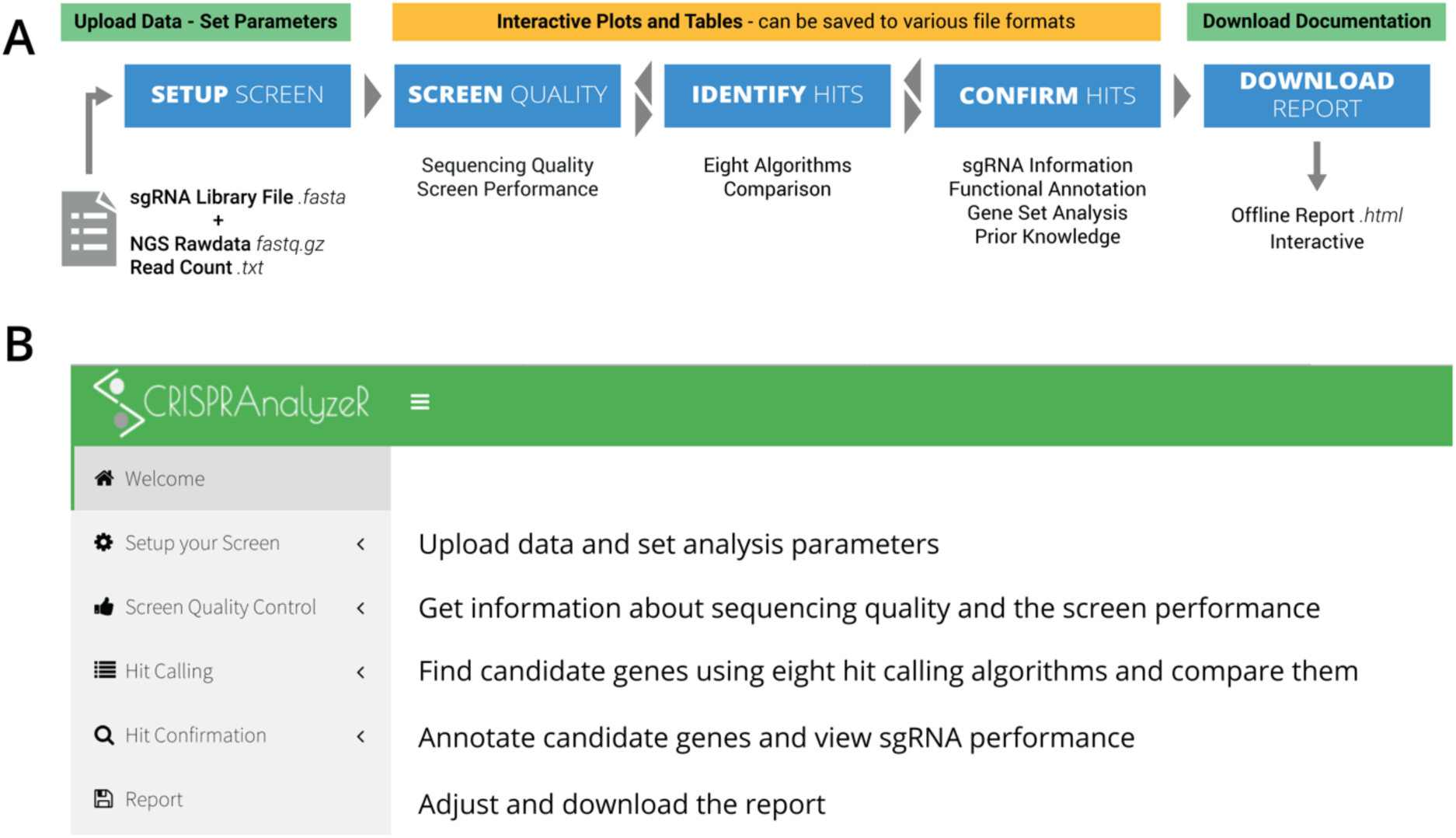
CRISPRAnalyzeR provides a user-friendly, end-to-end workflow for data analysis and documentation. **A**| The CRISPRAnalyzeR data analysis workflow is easy-to-use and only requires the upload of a sgRNA library file in addition to either sequencing or read count data. Interactive plots and tables are present throughout all steps. At the final stage, an interactive report can be downloaded with all plots and data included. **B**| The CRISPRAnalyzeR offers a user interface that is accessed via a web browser.

CRISPRAnalyzeR only requires the upload of two different file types: 1) an sgRNA library file in FASTA format which describes the applied sgRNA library and 2) screening data either as read count files (.txt) or as raw data sequencing files (.fastq.gz). For convenience, we provide pre-configured sgRNA library files for the most common sgRNA libraries available from Addgene (http://www.addgene.org.) We also provide sample data from a 12k sgRNA screen for resistance against TNF-related apoptosis-inducing ligand (TRAIL) (Heigwer *et al.*, 2016; Johnstone *et al.*, 2008). Moreover, data from a previously published genome-wide cell viability screen (Steinhart,Z. *et al.*, 2016) is provided. New users can analyze these example data sets to get a first impression of the software. After uploading the input data, the user sets the screening and analysis parameters (confidence intervals, controls, treatment groups, read count thresholds and gene identifiers) and starts the data analysis process. While hit-calling algorithms are executed, the user can browse an overview of all uploaded data and mapping statistics. They can download a FASTQ quality report as well as read count files. Once hit calling has been performed, users can assess screen performance, identify and annotate candidate genes and download an interactive report to document screen and analysis.

## Assessment of Sequencing Quality and Screen Performance

Ensuring data quality is an important premise to guarantee robust downstream analysis. CRISPRAnalyzeR initially guides users to the ‘Screen Quality Control’ section. This part of the application is divided into six panels for a comprehensive overview. These contain plots and tables ordered into logical groups that address similar aspects of quality control. To verify high sequencing quality, read width, read frequency as well as cycle-specific GC content are visualized. Additional statistics regarding base call proportion and average quality scores, determined using the RQC package (Souza and Carvalho, 2016) complement these information. The quality of the screen can be analyzed in the `Screen Read Count` panel. Here, information about the reproducibility of replicates, read depth, read count distributions (Figure 3A), controls as well as a series of statistical parameters (median, mean, min, max read count) are displayed for each sample. Additional visualizations in the `sgRNA Coverage` panel illustrate missing sgRNAs or genes as well as missing control sgRNAs and provide information about a loss in sgRNA coverage. User can examine in detail the replicate quality on both sgRNA and gene read count levels using a scatter matrix. Samples can be compared in the `Sample Comparison` panel based on their read counts or fold changes. Additional interactive scatter plots highlight any gene/sgRNA present in the screen including non-targeting and positive controls. As an example, Figure 3A shows a scatter plot where positive control sgRNAs for CASP8 (Caspase 8) are highlighted in red and sgRNAs against TNFRSF10B (TRAIL receptor 2) are in orange. Both CASP8 and TNFRSF10B are important mediators in the TRAIL pathway (Johnstone *et al.*, 2008). Finally, a principal component analysis (Ringnér, 2008) plot and clustered heat maps of read count data complete the quality control section. By offering a broad set of visualizations and statistics in this section, we aim to encourage users to confirm data quality for a robust downstream analysis.

**Figure 2:**
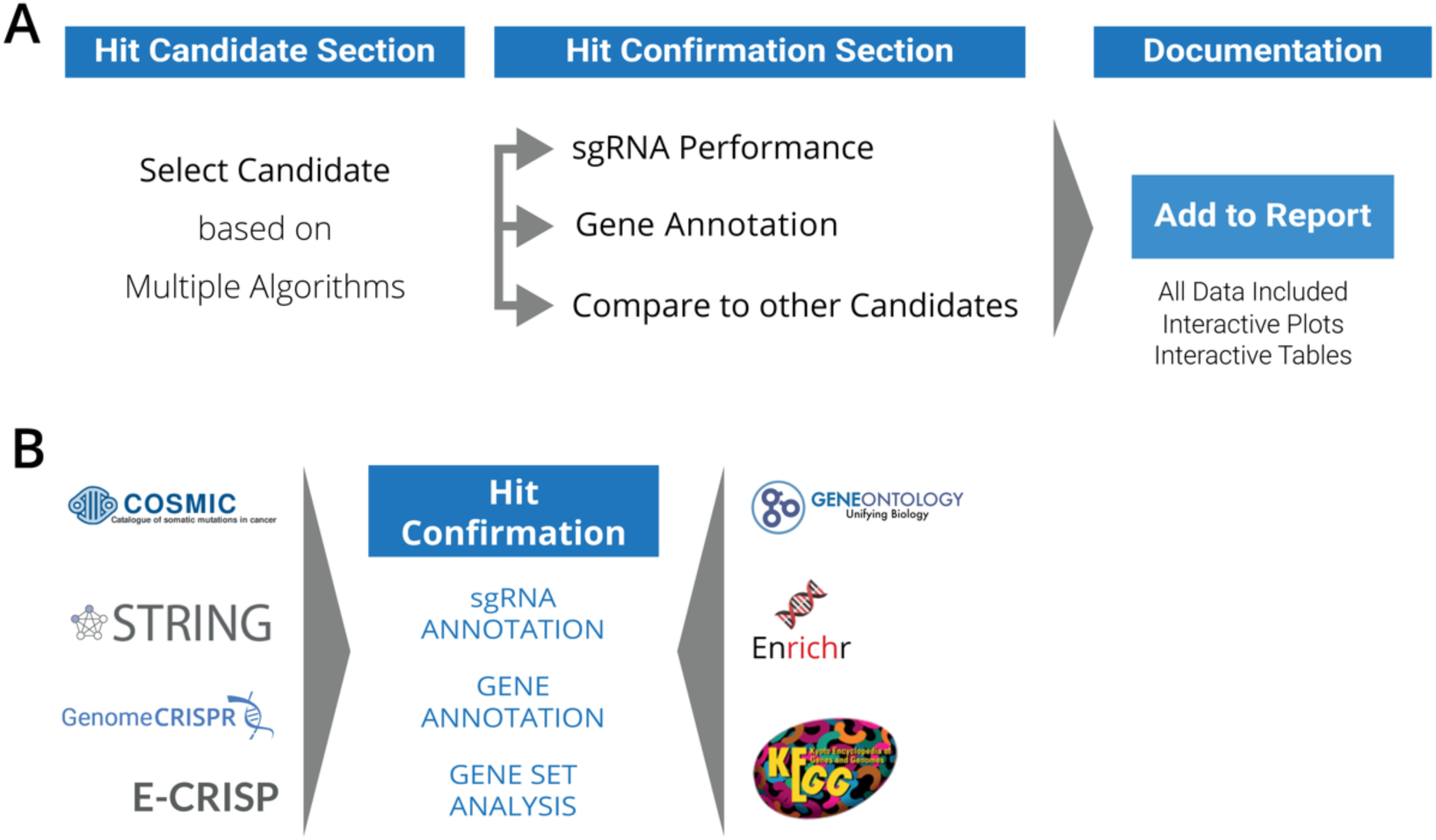
Annotation of candidate genes with additional information in a single step process. **A**| Based on the analysis output of multiple analysis algorithms, the user can select a candidategene, have a detailed look on the sgRNA performance, obtain gene annotations and compare the candidate gene with up to 20 other genes. All information about the candidate gene can be stored in the report for detailed documentation. **B**| In the `Hit Confirmation section`, users find information about all sgRNAs for a selected gene, obtain additional information from up to 26 data resources and perform a gene set analysis.

**Figure 3:**
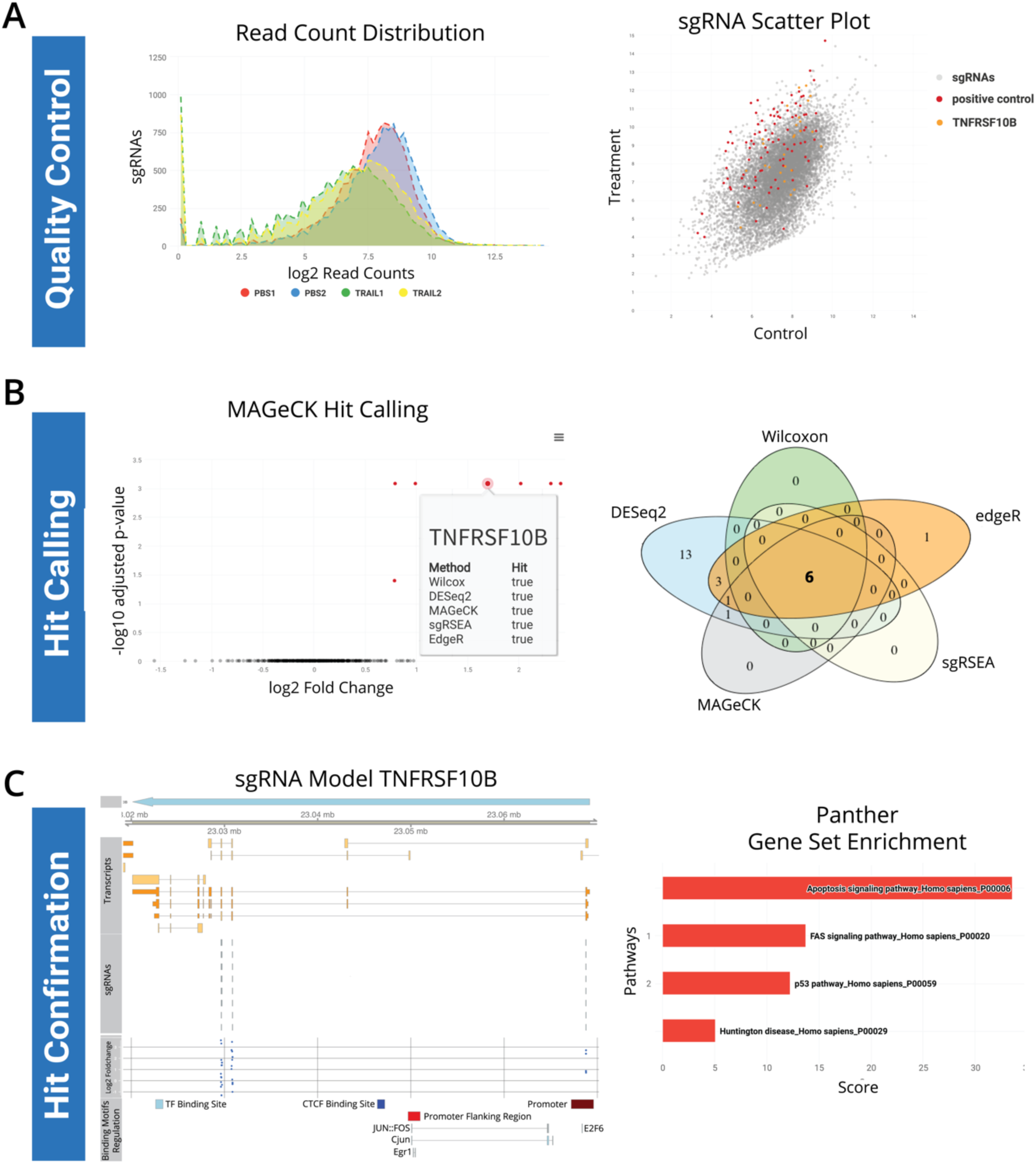
Examples of implemented result visualizations. **A**| In the ‘Screen Quality Control’ section interactive plots such as read count distributions and adjustable scatter plots provide an idea about the overall screen and replicate quality. **B**| For each analysis algorithm, the user is provided with visualizations regarding the fold change and adjusted p-values, allowing a direct comparison to the other algorithms. A VENN diagram further illustrates overlaps in candidate genes. **C**| In the `Hit Confirmation Section`, sgRNA and gene model visualizations link the sgRNA phenotype with its genomic location. Without leaving the application, a gene set analysis can be used to retrieve common information about up to 20 genes simultaneously.

## Identification of Candidates using Multiple Analysis Methods

Perhaps the most important step is the identification of candidate genes (‘hits’) for follow-up experiments. CRISPRAnalyzeR dedicates an entire section to this process where users can leverage eight different analysis algorithms to examine gene enrichment, depletion and gene essentiality between two treatment groups (Table 1). Per default, all eight analysis algorithms run simultaneously, allowing users to compare in detail the results retrieved by each individual method. This is a key feature to identify robust hits as the performance of each algorithm varies depending on data quality and experimental setup. To ensure a consistent data analysis, data is normalized using the DESeq2 size-factor estimation and variance stabilization as implemented in the DESeq2 package for R/Bioconductor (Love *et al.*, 2014). Then, hit calling is performed using both modern algorithms DESeq2 (Love *et al.*, 2014), MAGeCK (Li *et al.*, 2014), edgeR (Robinson *et al.*, 2010; Dai *et al.*, 2014), sgRSEA (Noh and Beibei, 2015), BAGEL (Hart and Moffat, 2016) and ScreenBEAM (Yu *et al.*, 2016) as well as classic statistical methods such as Z-score transformation (Cheadle *et al.*, 2003) or a Mann-Whitney hypothesis test. CRISPRAnalyzeR visualizes the results of the hit calling process, displaying p-value and fold change distributions as well as gene ranking plots. Furthermore, a detailed list of all hits is provided as a table that can be exported in various formats (Figure 3B). Finally, VENN diagrams illustrate overlaps of hits among different methods.

## Annotation of Candidate Genes

After the identification of candidate genes, it can be important to add additional annotation information based on public database. These annotations can form first biological hypotheses to guide further experimentals. CRISPRAnalyzeR contains a ‘Hit Confirmation’ section that serves this purpose. It is structured into five panels, where users can take advantage of data from 26 external resources (Figure 2B, Supplemental Table 2), allowing them to put candidate genes into biological context. First of all, CRISPRAnalyzeR evaluates each sgRNA using E-CRISP (Heigwer *et al.*, 2014). The results are visualized in the ‘sgRNA performance’ panel where reagent phenotypes such as read counts, fold changes, efficiency scores (Heigwer *et al.*, 2014) and predicted genomic binding sites are displayed. Here, users can in detail examine statistics for each individual sgRNA, including reproducibility across replicates. Next, the ‘Compare Genes’ panel examines in contrast sgRNA performance of up to 20 genes allowing for a direct comparison of candidate genes. More general information about a gene of interest is displayed in the ‘Gene Overview’ panel. These include phenotypes from previously published CRISPR screens reported in GenomeCRISPR (Rauscher *et al.*, 2017), Gene Ontology (Gene Ontology Consortium, 2015) annotations or associated KEGG pathways (Kanehisa *et al.*, 2017). A gene model view visualizes sgRNA target sites (Figure 2C). In addition, CRISPRAnalyzeR includes data derived from ENSEMBL (Durinck *et al.*, 2005; Aken *et al.*, 2016), COSMIC (Forbes *et al.*, 2015) and Enrichr (Kuleshov *et al.*, 2016) including predicted miRNA binding sites, protein interactions/motifs and somatic mutations in cancer cells. Within CRISPRAnalyzeR, users can perform gene set analysis based on 17 different data resources using the Enrichr web service (Chen *et al.*, 2013; Kuleshov *et al.*, 2016) (Figure 2C) or enrich genes with up to 200 annotations from ENSEMBL. All results can be added to the analysis report (Figure 2A) allowing for streamlined documentation of results.

## Screening Documentation Using the Interactive Offline Report

CRISPRAnalyzeR documents in detail all analysis steps performed, including generated data, plots, tables and annotations. A printable report can be exported in which the core information is included per default. Additionally, users can add any additional plot on demand per click on the ‘Add to report’ buttons. The report can be downloaded as an HTML file. It is based on R bookdown (Xie, 2016) and is fully interactive, supporting a full-text search. The ‘Report Section’ provides options for users to configure the contents of the report. Providing automated documentation of all analyses we aim to promote transparent and reproducible data science.

## Availability of the CRISPRAnalyzeR

CRISPRAnalyzeR ((http://www.crispr-analyzer.org) can be downloaded from GitHub as source code and pre-configured application for a platform-independent use (http://www.github.com/boutroslab/CRISPRAnalyzeR.). Furthermore, sample data as well as pre-generated sgRNA library files can be obtained from within the web-tool or from the GitHub page. The availability of tutorials in video format and additional help on the GitHub page or in the respective sections of the web-tool assist the user. Users are encouraged to try our online live demonstration available at http://www.crispr-analyzer.org.

## Acknowledgements

We thank Enrico Girardi and Ulrich Goldmann (CeMM Forschungszentrum für Molekulare Medizin, Vienna), Kwang Lee and Arek Kendirli for testing and feedback.

## Funding

This work was in part supported by an ERC grant.

## References

Cheadle, C. et al. (2003) Analysis of microarray data using Z score transformation. J. Mol. Diagn., 5, 73–81.

Chen, E.Y. et al. (2013) Enrichr: interactive and collaborative HTML5 gene list enrichment analysis tool. BMC Bioinformatics, 14, 128.

Dai, Z. et al. (2014) edgeR: a versatile tool for the analysis of shRNA-seq and CRISPR-Cas9 genetic screens. F1000Research, 3, 95.

Diaz, A.A. et al. (2014) HiTSelect: a comprehensive tool for high-complexity-pooled screen analysis. Nucleic Acids Res.

Gene Ontology Consortium (2015) Gene Ontology Consortium: going forward. Nucleic Acids Res., 43, D1049–56.

Hart, T. et al. (2015) High-Resolution CRISPR Screens Reveal Fitness Genes and Genotype-Specific Cancer Liabilities. Cell, 163, 1515–1526.

Hart, T. and Moffat, J. (2016) BAGEL: a computational framework for identifying essential genes from pooled library screens. BMC Bioinformatics, 17, 164.

Heigwer, F. et al. (2016) CRISPR library designer (CLD): Software for multispecies design of single guide RNA libraries. Genome Biol., 17.

Heigwer, F. et al. (2014) E-CRISP: fast CRISPR target site identification. Nat. Methods, 11, 122–123.

Johnstone, R.W. et al. (2008) The TRAIL apoptotic pathway in cancer onset, progression and therapy. Nat. Rev. Cancer, 8, 782–98.

Kanehisa, M. et al. (2017) KEGG: new perspectives on genomes, pathways, diseases and drugs. Nucleic Acids Res., 45, D353–D361.

Kuleshov, M. V et al. (2016) Enrichr: a comprehensive gene set enrichment analysis web server 2016 update. Nucleic Acids Res., 44, W90–7.

Li, W. et al. (2014) MAGeCK enables robust identification of essential genes from genome-scale CRISPR/Cas9 knockout screens. Genome Biol., 15, 554.

Li, W. et al. (2015) Quality control, modeling, and visualization of CRISPR screens with MAGeCK-VISPR. Genome Biol., 16, 281.

Love, M.I. et al. (2014) Moderated estimation of fold change and dispersion for RNA-seq data with DESeq2. Genome Biol., 15, 550.

Noh, J. and Beibei, C. (2015) sgRSEA: Enrichment Analysis of CRISPR/Cas9 Knockout Screen Data.

Rauscher, B. et al. (2017) GenomeCRISPR - a database for high-throughput CRISPR/Cas9 screens. Nucleic Acids Res., 45, D679–D686.

Ringnér, M. (2008) What is principal component analysis? Nat. Biotechnol., 26, 303–304.

Robinson, M.D. et al. (2010) edgeR: a Bioconductor package for differential expression analysis of digital gene expression data. Bioinformatics, 26, 139–140.

Shalem, O. et al. (2014) Genome-scale CRISPR-Cas9 knockout screening in human cells. Science, 343, 84–7.

Shi, J. et al. (2015) Discovery of cancer drug targets by CRISPR-Cas9 screening of protein domains. Nat. Biotechnol., 33, 661–667.

Souza, W. and Carvalho, B. (2016) Rqc: Quality Control Tool for High-Throughput Sequencing Data.

Steinhart, Z. et al. (2016) Genome-wide CRISPR screens reveal a Wnt-FZD5 signaling circuit as a druggable vulnerability of RNF43-mutant pancreatic tumors. Nat. Med.

Wang, T. et al. (2015) Identification and characterization of essential genes in the human genome. Science (80-.)., 350, 1096–1101.

Winter, J. et al. (2015) caRpools: An R package for exploratory data analysis and documentation of pooled CRISPR/Cas9 screens. Bioinformatics.

Xie, Y. (2016) Authoring Books and Technical Documents with R Markdown https://bookdown.org.

Yu, J. et al. (2016) ScreenBEAM: a novel meta-analysis algorithm for functional genomics screens via Bayesian hierarchical modeling. Bioinformatics, 32, 260–7.

Zhou, Y. et al. (2014) High-throughput screening of a CRISPR/Cas9 library for functional genomics in human cells. Nature, 509, 487–91.

